# Efficacy and safety of an mRNA-based RSV preF vaccine in preclinical studies

**DOI:** 10.1101/2024.09.29.615646

**Authors:** Huarong Bai, Qin Li, Yue Gao, Yubin Zhao, Xueliang Yu, Rongkuan Hu

## Abstract

The transmembrane fusion (F) protein of RSV plays important roles in RSV pathogenesis as it mediates the fusion between virus and the target cell membrane. During the fusion process, F protein transits from a metastable state (prefusion, preF) to a stable state after merging of virus and cell membranes (postfusion, postF). The majority of highly neutralizing antibodies induced by natural infection or immunization targets the preF form, making it the preferred antigen for vaccine development. Here, we evaluate the mRNA vaccine candidate, STR-V003, which encapsulates the modified mRNA encoding the preF protein in lipid nanoparticles (LNPs). This vaccine demonstrated robust immunogenic in both mice and cotton rats. STR-V003 induced high levels of neutralizing antibodies and RSV preF-specific IgG antibodies, and significantly reduced the RSV viral loads in the lung and nose tissue of challenged animals. In addition, STR-V003 did not have obvious enhancement of lung pathology without causing vaccine enhanced disease (VED). The repeated dose general toxicology studies and local tolerance studies of STR-V003 were evaluated in rats. Therefore, STR-V003 has an acceptable safety profile and robust protective immunity against RSV, and has been approved by the FDA to enter phase I clinical study (NCT06344975).

## Introduction

Human respiratory syncytial virus (RSV) is an enveloped, single-stranded, and negative-sense RNA virus which causes a major burden in public health, both in developing and in industrialized countries. There are two RSV sub-types, RSV A and RSV B, which both cocirculate and cause illnesses during each respiratory season.[1, 2]

The transmembrane fusion (F) protein of RSV plays important roles in RSV pathogenesis as it mediates the fusion between virus and the target cell membrane. During the fusion process, F protein transits from a metastable state (prefusion, preF) to a stable state after merging of virus and cell membranes (postfusion, postF)[3]. The majority of highly neutralizing antibodies induced by natural infection or immunization targets the preF form, making it the preferred antigen for vaccine development.[4-7] F protein contains antigenic determinants associated with neutralizing antibodies and cytotoxic T-lymphocyte (CTL)-mediated immunity. Some antigenic sites (II and IV) are common between both pre- and post-fusion forms, while others are form-specific (e.g., □ and Φ are preF specific).[1, 8, 9]

To date, two RSV protein subunit vaccines which induces immune responses against RSV preF protein were approved by FDA for adults 60 years old and over, namely, Arexvy (developed by GSK) and Abrysvo (developed by Pfizer).[10] The latter vaccine (Abrysvo) was also approved by FDA for pregnant women to protect newborns and infants against severe RSV disease in the first 6 months after birth.[11]. Moderna’s mRNA-based mResvia became the third RSV vaccine approved by the FDA.[12]STR-V003 is an mRNA vaccine produced by encapsulating the mRNA that encodes the prefusion F protein (preF) of respiratory syncytial virus (RSV) in lipid nanoparticles (LNPs). There are six codon mutations in the mRNA sequence to stabilize the prefusion form of the F protein and increase the preF protein expression. Here, we demonstrated that STR-V003 was able to induce the RSV preF-specific IgG antibodies and neutralizing antibodies against the RSV in mice and cotton rats. Furthermore, STR-V003 vaccine could significantly reduce the RSV viral loads in the lung and nose tissue of challenged animals. Meanwhile, STR-V003 did not cause any obvious enhancement of lung pathology. In addition, the repeated dose general toxicology studies and local tolerance studies of STR-V003 were evaluated in rats. Collectively, the results showed that STR-V003 had an acceptable safety profile and provided efficient protection against RSV infection without causing vaccine enhanced disease (VED) in both mouse and cotton rat RSV infection models, suggest that STR-V003 is a potential vaccine candidate to prevent RSV infectious diseases and has been approved by the FDA IND.

## Results

### Lipid Synthesis and Screening of LNP formulations

The asymmetric alkanolamines ionizable lipids (AAiL) library was crafted through a precise Michael addition process between amine alcohols and acrylates through rational design (Fig. 1a, 1b). These AAiL synthetic products were then formulated with DSPC, cholesterol, DMG-PEG2000, and luciferase (Luc) mRNA to create LNPs for an initial assessment with a fixed molar ratio of (50:10:38.5:1.5) for initial screening (Fig. 1c, 1d). The lipid of A1-EP10-O18A was identified for further optimization using a rigorous Design of Experiments (DOE) strategy (Fig. 1e, Supplementary Figure 1 and Supplementary Figure 2). The refined F11 formulation packing with luciferase mRNA (STAR0225) was chosen to compared with Moderna’s formulation (SM102) and showed a better local distribution. (Fig. 1f, 1g).

**Fig. 1.**
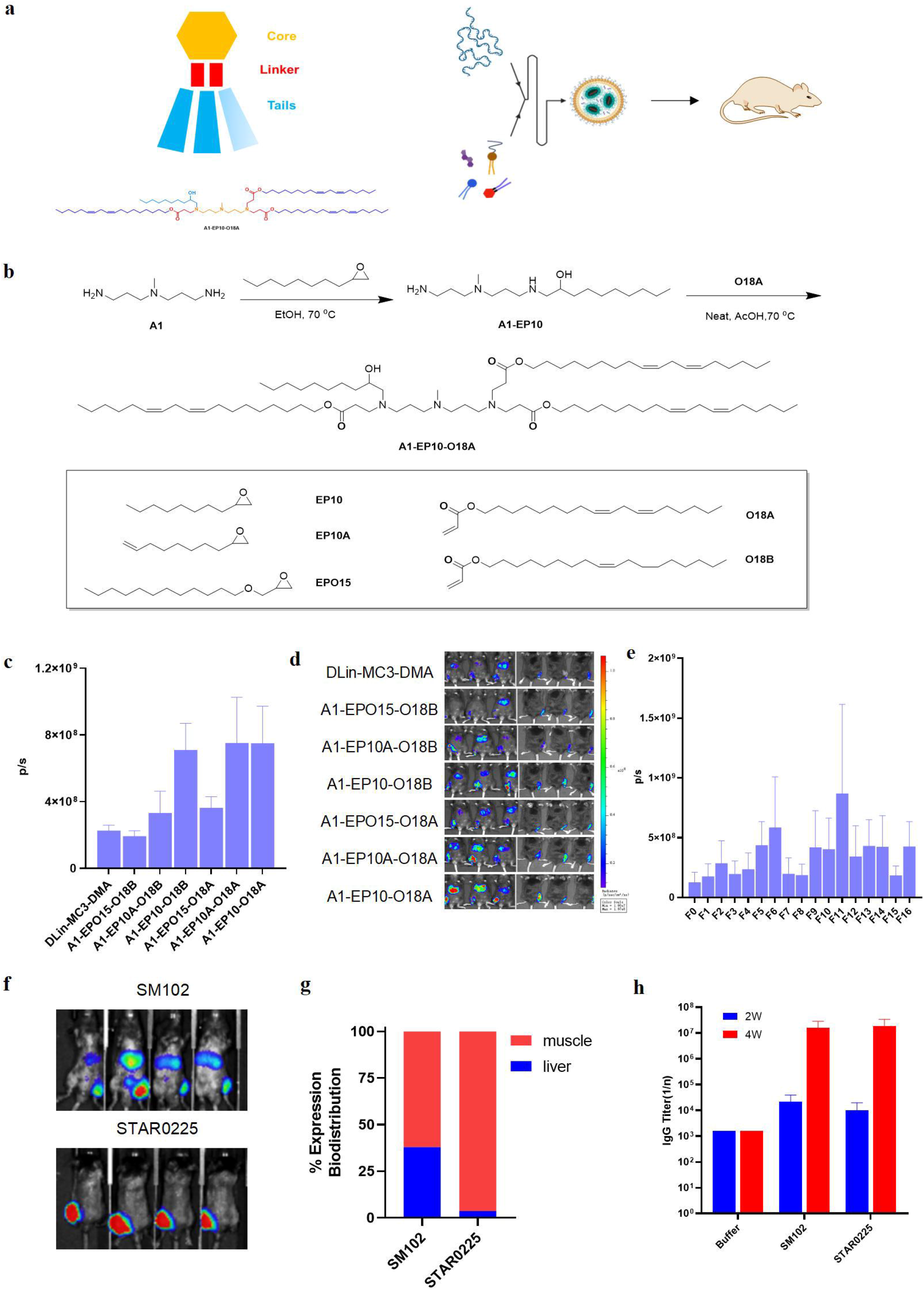
Design, Screening and Optimization of LNP Formulation. **(a)** Schematic of AAiL structure and LNPs formulating for mRNA delivery. **(b)** Structure of tails and synthetic route toward A1-EP10-O18A. **(c)** In vivo delivery efficacy of luciferase mRNA in C57BL/6 mice (0.25 mg/kg mRNA per mouse, through I.M. injection, n = 3, mean ± SEM. Luminescence were recorded 5 h after injection. **(d)** Imaging of body were recorded 5 h after injection. **(e)** DOE study results of in vivo delivery efficacy of luciferase mRNA in C57BL/6 mice (0.25 mg/kg mRNA per mouse, through I.M. injection, n = 2, mean ± SEM. Luminescence were recorded 5 h after injection. **(f, g)** Comparison of Luminescence of STAR0225 and SM102 LNP in C57BL/6 mice (0.25 mg/kg through I.M. injection); **(h)** Comparison of wild type RSV F protein encapsulation with STAR0225 and SM102 LNP in BALB/c mice (5 μg/mice through I.M. injection)

**Fig. 2:**
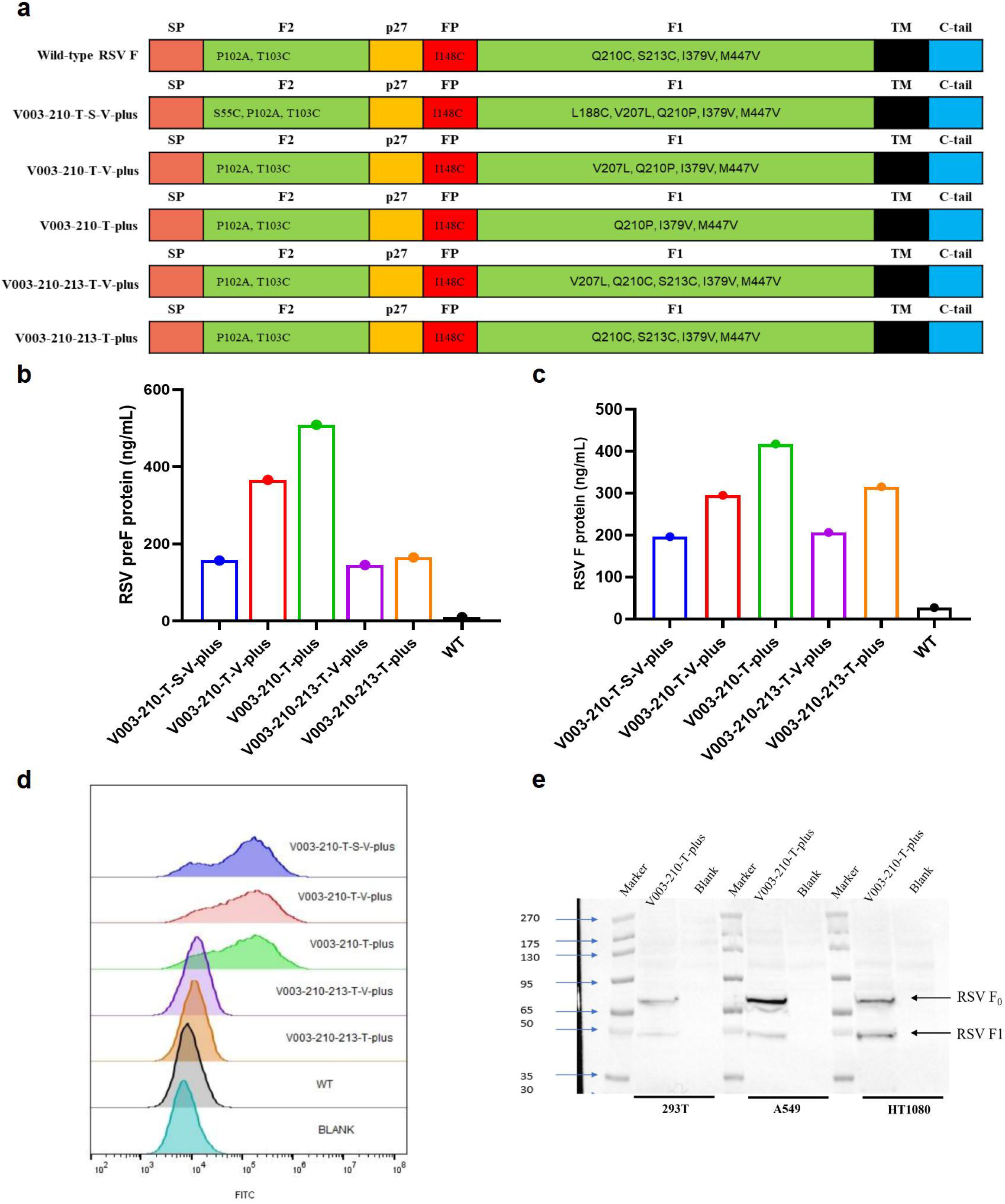
Sequence screening for STR-V003 based on in vitro expression and characterization of preF protein in cell lines. **(a)** RSV F proteins encoded by different mRNA constructs. Wild-type (WT) RSV F encoding the F protein of the prototype RSV A2 strain. SP, signal peptide; F2, a polypeptide of F protein; p27, p27 peptide; FP, fusion peptide; F1, a polypeptide of F protein; TM, transmembrane peptide; C-tail, cytoplasmic tail; point mutations are labeled. **(b)** Expression level of RSV preF proteins in 293T cells. (**c)** Expression level of RSV F proteins in 293T cells. **(d)** Expression of RSV preF proteins in HT1080 cells detected by FACS. Fluorescent (Alexa Fluor 488)-labeled antibody recognizing antigenic site Φ (specific for RSV preF protein) was used to stain RSV preF protein. **(e)** Maturation of RSV F protein. F_0_, inactive precursor of F protein.

Furthermore, we investigated the immunogenicity of mRNA, coding wild-type (WT) F protein of the RSV A2 strain, encapsulated in SM102 and STAR0225 in mice. WT-LNPs (both at 10μg) were used to immunize (IM injections, at Day 0 and Day 21) mice (5/female/group). Serum samples were collected on days 14 and 28. The RSV F-specific antibody IgG titers were analyzed 2 and 4 weeks after the first dose by ELISA. As shown in Figure. 1h, WT-SM102 and WT-STAR0225 induced comparable RSV F-specific antibody IgG titers.

### In vitro screening of RSV preF mRNA sequence for STR-V003

The majority of highly neutralizing antibodies induced by natural infection or immunization targets the RSV preF form, making it the preferred antigen for vaccine development. STR-V003 is an mRNA vaccine produced by encapsulating the mRNA that encodes the preF of respiratory syncytial virus RSV in LNPs. The mRNA molecule was designed according to the wild-type (WT) F protein sequence of the prototype RSV A2 strain. Following methods used formerly for SARS-CoV-2 S protein and RSV F protein, such as engineered disulfide mutations, electrostatic mutations, cavity filling mutations and trimerization domain[5, 13-15], we constructed F protein mutants to increase expression yields of F and preF protein by the introduction of the amino acid mutations including amino acid substitutions, deletions, or additions relative to an RSV WT F protein in our earlier studies(PCT/CN2024/083234). Combined with the findings in the in vitro expression study, we finally selected five mutant mRNAs for subsequent studies (Fig. 2a).

To optimize the mRNA sequence used in the vaccine, five mRNA constructs encoding RSV F and preF proteins were transfected into HT1080, A549, or 293T cells. The expression levels of RSV F and preF proteins were analyzed by ELISA and flow cytometry (FACS) in cells transfected with different mRNAs. As shown in Fig. 2b, Fig. 2c and Fig. 2d, compared with V003-WT, all mutant mRNAs led to higher expression levels of F and preF protein 24 hrs after transfection. Taken together, V003-210-T-plus is among the mRNA constructs that generally yielded the highest expression under different experiment conditions. Moreover, RSV F protein specific antigenic sites of V003-210-T-plus were detected by FACS (Supplementary Figure 3). Except for antigenic site I, all other antigenic sites could be identified. It might be possible for antibody against antigenic site I binding tighter to the post-F conformation than pre-F conformation.

The inactive precursor, RSV F_0_, requires cleavage during intracellular maturation by a furin-like protease.[16, 17] RSV F contains two furin sites, which leads to three polypeptides: F2, p27(a 27-amino acid fragment) and F1.[18-20] The cleavage process was analyzed by Western blot in HT1080, A549, or 293T cells transfected with mRNA V003-210-T-plus. As shown in Fig. 2e, two forms of RSV F proteins were detected, which migrate at ∼65kDa (RSV F_0_) and ∼50 kDa (RSV F1). The detection of the faster-migrating variant of RSV F protein confirmed the cleavage of inactive precursor (RSV F_0_).

### In vivo screening of RSV preF mRNA sequence for STR-V003 in mice and rats

The candidate mRNA sequences were screened based on the ability to induce RSV F or preF protein-specific IgG in BALB/c mice and Sprague Dawley (SD) rats. RSV preF mRNAs with different sequence optimization strategies were produced by in vitro transcription and encapsulated in Starna’s LNPs STAR0225. The LNPs (in mice, 50 µl, containing 0, 5, or 10 µg mRNA; in rats, 500 µl, containing 50 µg mRNA) were used to immunize mice or SD rats (3/sex/group) via intramuscular (IM) injections on Day 0 and Day 14. Body weight was measured every 4 days until the end of the study. The RSV preF or RSV F-binding antibody IgG titers were analyzed 2 and 5 weeks after the first dose by ELISA.

No significant difference in body weight was observed between the vehicle group and dosing groups. As shown in Fig. 3a and Fig. 3b, compared with the V003-WT, all mutant mRNAs induced higher RSV F and RSV preF-binding antibody IgG titers in mice at both 2 weeks and 5 weeks after the first dose. Compared with the 1st dose in mice, the 2nd dose of vaccine led to higher titers of RSV F and RSV preF-specific antibody. As shown in Fig. 3c and Fig. 3d, compared with the V003-WT, all mutant mRNAs induced higher RSV F-binding antibody IgG titers in rats at both 2 weeks and 5 weeks after the first dose. Compared with the 1st dose in rats, the 2nd dose of vaccine induced higher RSV F-binding antibody IgG titers, demonstrating that 2nd dose of vaccine further boosted the immune response.

**Fig. 3:**
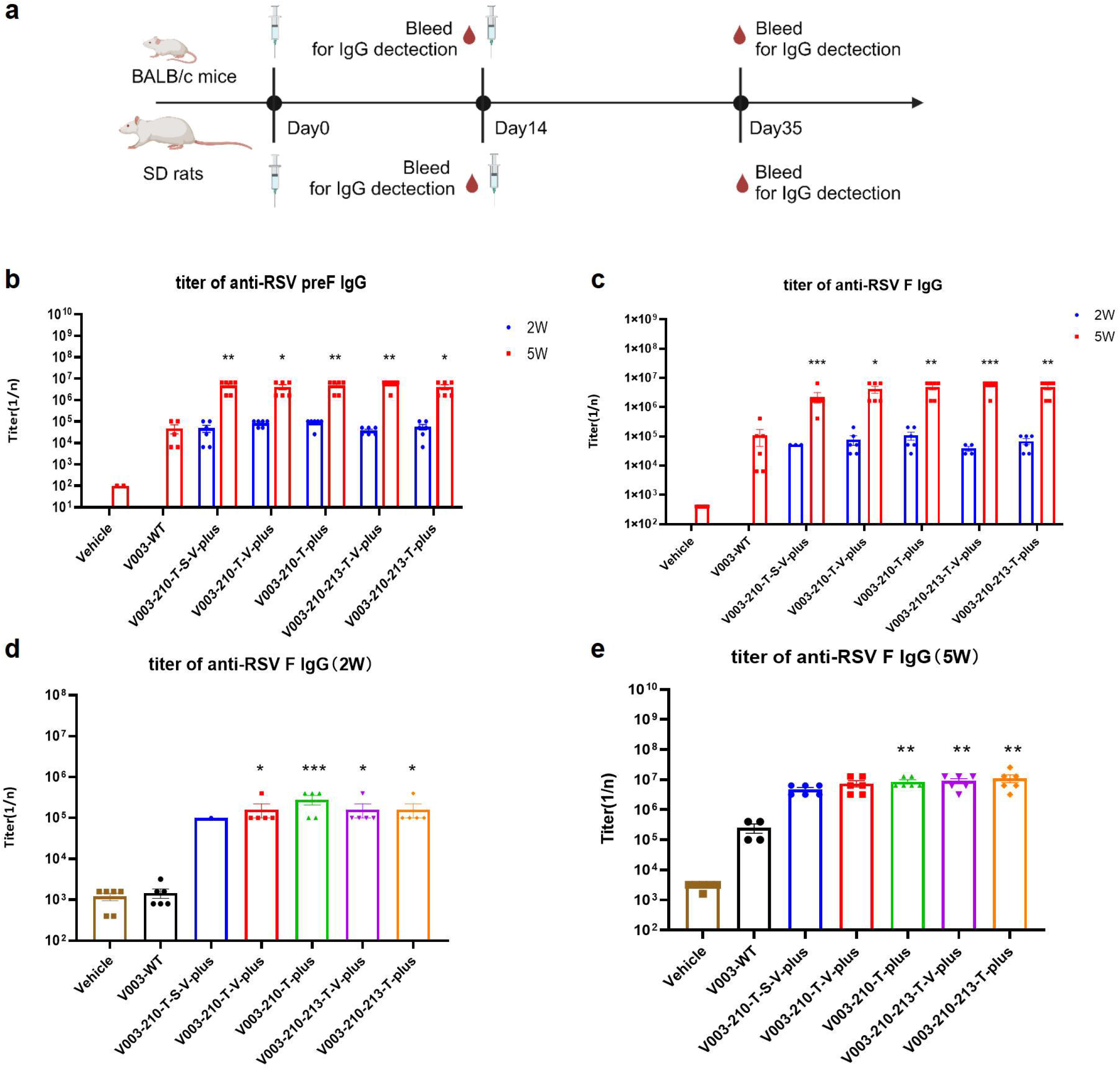
Optimization of antigen mRNA sequence in BALB/c mice and SD rats. **(a)** The schematic diagram of the immunogenicity of STR-V003 in mouse and rat model. RSV F mRNAs with different sequence encapsulated in LNPs. The LNPs (in BALB/c mice, at 50 µL, containing 5µg mRNA, in SD rats, at 500µL, containing 50µg mRNA) were used to immunize mice (3/sex/group) and SD rats (3/sex/group) via IM injections on Day 0 and 2 weeks. The RSV preF/F-binding antibody IgG titers were analyzed 2 and 5 weeks after the first dose by ELISA. **(b, c)** Titers of RSV preF/F-binding antibodies in mice at 2 and 5 weeks. **(d, e)** Titers of RSV F-binding antibodies in rats at 2 and 5 weeks. Data is shown as mean ± SEM and Asterisks indicate significant differences. (*: p<0.05, **: p<0.01, ***: p<0.001, compared with V003-WT at 2W or 5W).

Taken together, V003-210-T-plus generally induced the highest RSV preF-binding antibody titers and RSV F-binding antibody titers. Combined with the findings in the in vitro expression study, these results suggested that V003-210-T-plus is the optimal mRNA sequence that can be used in the STR-V003 vaccine.

### Comparison of immunogenicity of STR-V003 and mRNA-1345 in mice

Moderna’s mRNA-based mResvia, mRNA-1345, was the third RSV vaccine approved by the FDA. We compared the immunogenicity of STR-V003 and mRNA-1345 in mice. STR-V003 and mRNA-1345 (both at 5μg) were used to immunize (IM injections, at Day 0 and Day 21) mice (5/female/group). Serum samples were collected on days 14, 28 and 35. The RSV F-specific antibody IgG titers were analyzed 2 and 4 weeks after the first dose by ELISA. The titer of neutralizing antibodies in serum samples of 5 weeks was detected by microneutralization. As shown in Fig. 4, STR-V003 and mRNA-1345 induced higher levels of RSV F-binding antibody IgG titers and neutralizing antibodies titers.

**Fig. 4:**
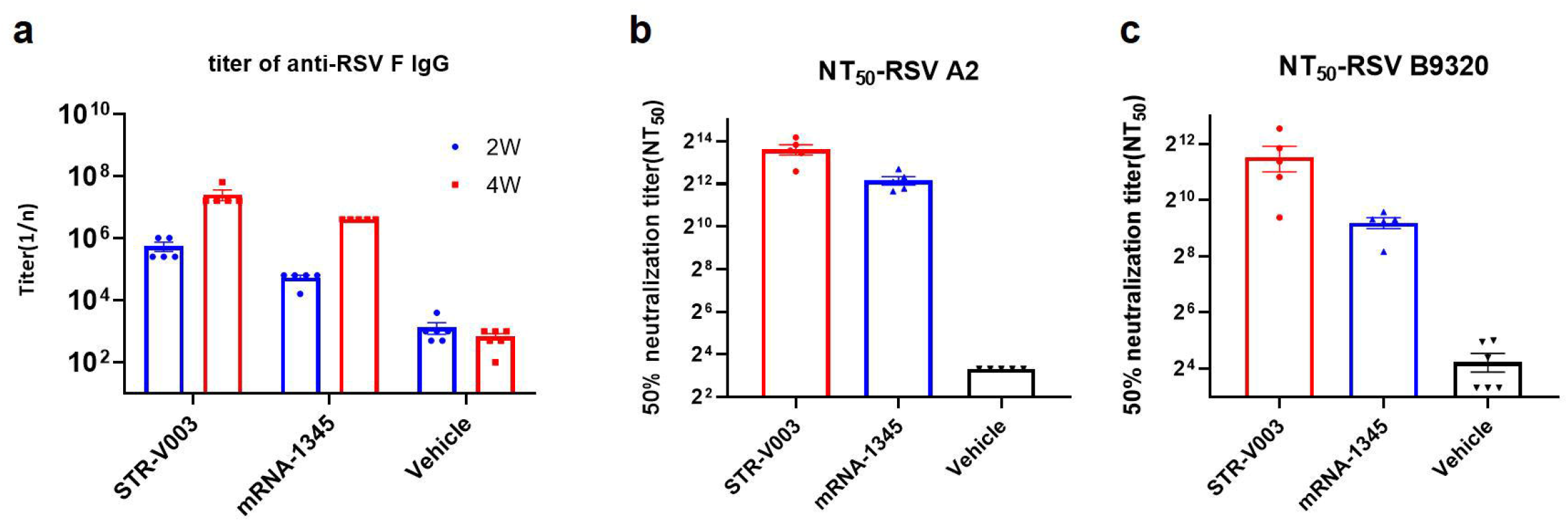
Comparison of immunogenicity of STR-V003 and mRNA-1345 in mice. STR-V003 and mRNA-1345(both at 5μg) were used to immunize (IM injections, at Day 0 and Day 21) mice (5/sex/group). Serum samples were collected on days 14, 28 and 35. (**a**) RSV F-specific IgG antibody in mouse serum on day14 and 28. **(b)** and **(c)** The 50% neutralizing titer (NT_50_) against RSV/A2 and RSV/B9320 in mice on day 35.

### Optimization of administration route, dose-response assessment and immunization regimen in BALB/c Mice

To optimize the route of immunization, V003-210-T-plus encapsulated in LNPs (STR-V003) were used to immunize mice (5/sex/group) via different routes (IM, subcutaneous (SC), or intradermal (ID) injections) on Day 0 and Day 21. Compared to the vehicle IM group, STR-V003 administered via different routes (IM, SC, and ID) all yielded similarly higher immunogenicity in mice (Fig. 5a). The most common clinical vaccination route, IM was selected for further studies.[21]

**Fig. 5:**
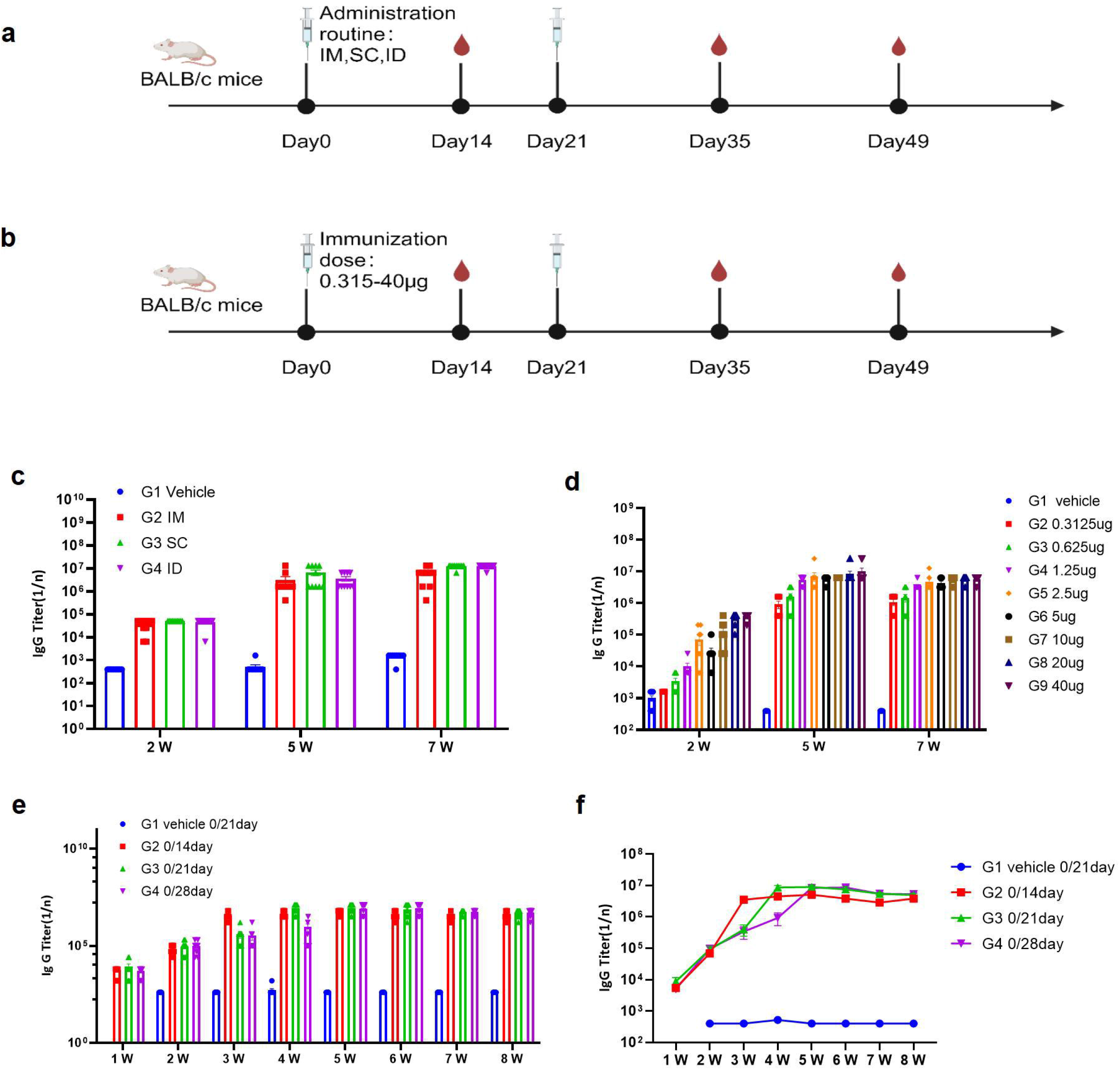
Optimization of administration route, dose response and immunization regimen in BALB/c mice. **(a)** Optimization of administration route in BALB/c mice. V003-210-T-plus mRNA was encapsulated in LNPs. The LNPs (50 µl, containing 5 µg mRNA) were used to immunize mice (5/sex/group) via different routes on Day 0 and Day 21. The immunogenicity was evaluated 2, 5 and 7 weeks after the first dose by measuring RSV F-binding antibody (IgG) titers. IM, intramuscular; SC, subcutaneous; ID, intradermal. **(b)** Optimization of dose response in BALB/c mice. The LNPs (approximately 50 µL, containing 0.3125, 0.625, 1.25, 2.5, 5, 10, 20, or 40 µg V003-210-T-plus mRNA) were used to immunize mice (5/sex/group) via IM injections on Day 0 and Day 21. Titers of RSV F-binding antibodies in mice at 2, 5 and 7 weeks were evaluated. **(c)** Titers of RSV F-binding antibodies in mice at 2, 5 and 7 weeks via different administration routes. **(d)** Titers of RSV F-binding antibodies in mice at 2, 5 and 7 weeks with different immunization dose. **(e, f)** The LNPs (50 µl, containing 5 µg V003-210-T-plus mRNA) were used to immunize mice (5/sex/group) via IM injections following different dosing regimens (Day 0 and Day 14, Day 0 and Day 21, Day 0 and Day 28). The immunogenicity was evaluated once weekly for up to 8 weeks after the first dose by measuring RSV F-binding antibody (IgG) titers.

To understand the effects of immunization doses on the induced immunogenicity, different doses of STR-V003 were used to immunize mice(5/sex/group) via IM injections on Day 0 and Day 21. At 2 weeks, RSV F-binding antibody IgG titers increased in a dose-dependent manner when the dose increased from 0.3125 to 40 μg. As shown in Fig. 5b, compared with those at 2 weeks, RSV F-binding antibody IgG titers increased at 5 weeks, suggesting that the 2nd dose of vaccine further boosted the immune response. RSV F-binding antibody IgG titers were not significantly different between 7 weeks and 5 weeks, suggesting that immune responses to the vaccine were maintained for at least 4 weeks after the 2nd vaccination. At both 5 and 7 weeks, RSV F-binding antibody IgG titers increased when the dose increased from 0.3125 to 1.25 μg but did not further increase when doses were higher than 1.25 μg.

To evaluate the impact of immunization regimen on the immunogenicity elicited, STR-V003 were used to immunize mice(5/sex/group) via IM injections following different dosing regimens (Day 0 and Day 14, Day 0 and Day 21, Day 0 and Day 28).

As shown in Fig. 5c and Fig. 5d, the 0/21-day regimen generated more potent immunogenicity than the 0/14-day regimen. Due to the earlier boost (3 weeks after the 1st dosing instead of 4 weeks), the 0/21-day regimen induced stronger immunogenicity than the 0/28-day regimen at 4 weeks after the 1st dosing. At other time points, the 0/21-day regimen induced comparable immunogenicity as the 0/28-day regimen. Taken together, the 0/21-day regimen is chosen for future trials.

### Immunogenicity and challenge protection of STR-V003 in mouse model

Mice and cotton rats, two commonly used models of human RSV infection, have provided insights into pathogenesis of RSV infections and mechanisms of immunity[22, 23]. The immunogenicity and challenge protection of STR-V003 were first evaluated in mouse model. The formalin inactivated vaccine (FI-RSV, at 0.05 μg), STR-V003 (at 1.25, 5, or 20 μg) was used to immunize (IM injections, at Day 0 and Day 21) mice (5/sex/group). RSV/A2 (ATCC, VR-1540) was inoculated intranasally (1.0×10^6^ PFU/animal) on Day 35 (Fig. 6a). Serum samples were collected on days 14, 28, and 35, and lung tissue samples were collected on day 40. The titer of neutralizing antibodies in serum samples was detected by microneutralization, the titer of RSV preF-specific IgG antibody in animal serum samples was detected by ELISA, the titer of RSV virus in lung tissue was detected by qPCR and the HE staining of lung tissue sections was evaluated for peribronchiolitis (PB), perivasculitis (PV), interstitial pneumonia (IP) and alveolitis (A),[5] and evaluate the immunogenicity and in vivo antiviral efficacy of the tested vaccine. Histopathology semiquantitative scoring criteria of lung tissue sections was shown in Supplementary Table 2.

**Fig. 6:**
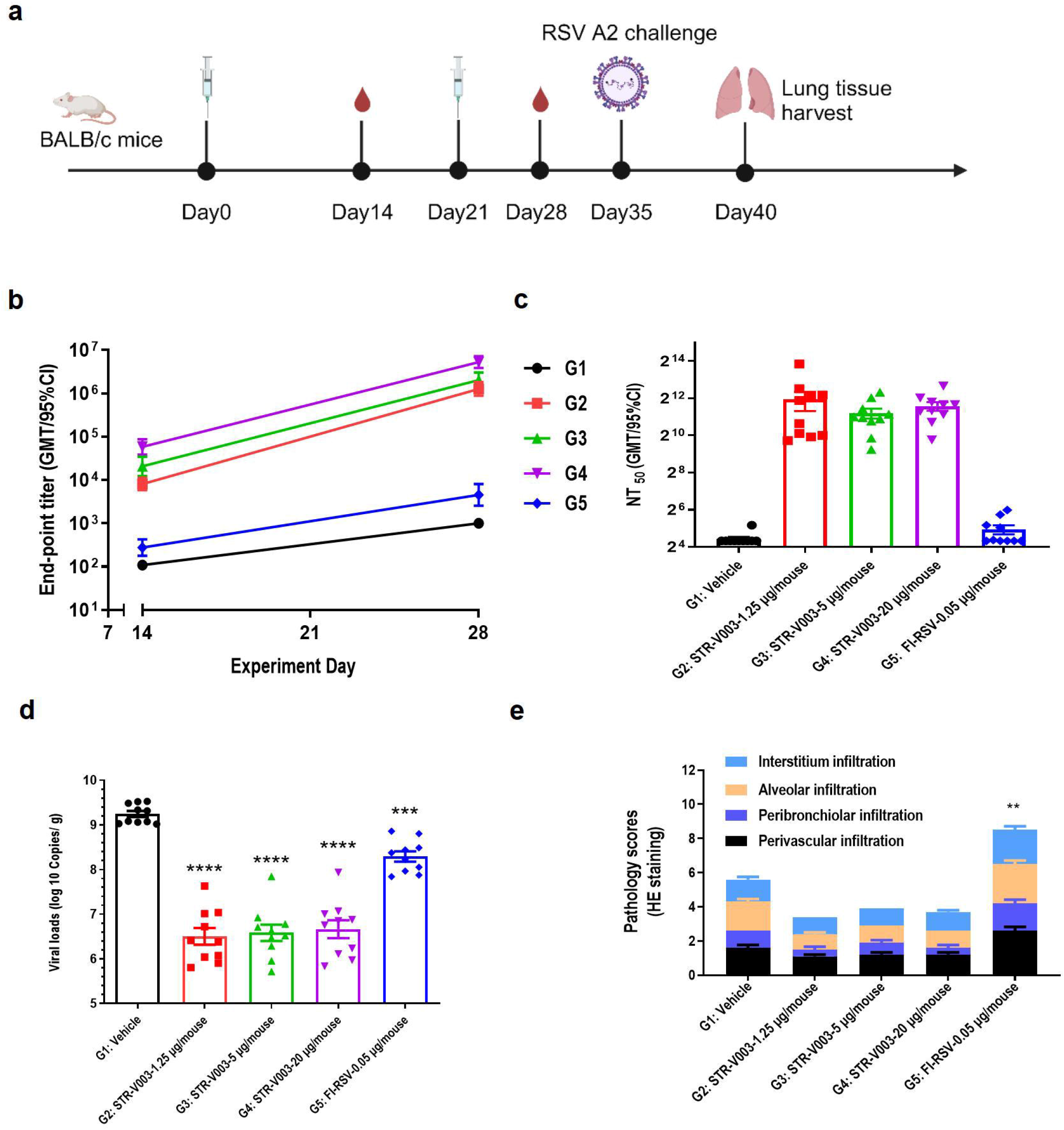
Protection Against RSV Challenge in Mouse Model. **(a)** The schematic diagram of the immunogenicity and challenge protection of STR-V003 in mouse model. The formalin inactivated vaccine (FI-RSV, at 0.05 μg), V003-210-T-plus encapsulated in LNPs (STR-V003, at 1.25, 5, or 20 μg) was used to immunize (IM injections, at Day 0 and Day 21) mice (5/sex/group). RSV/A2 (ATCC, VR-1540) was inoculated intranasally (1.0×10^6^ PFU/animal) on Day 35. Serum samples were collected on days 14, 28 and 35 and lung tissue samples were collected on day 40. (**b)** RSV preF-specific IgG antibody in mouse serum on day14 and 28. (**c)** The 50% neutralizing titer (NT_50_) in mice on day 35. (**d)** Viral loads in mouse lung tissue on day 40. (**e, f)** Pathology scores and H&E staining of mouse lung tissues on day 40.

During the study, no significant weight loss was observed. Compared with the vehicle, FI-RSV significantly reduced the RSV virus titer in the lung tissue by 0.96 Log10 (copies/g lung) (Fig. 6d). Total lung pathology score in FI-RSV group was 8.5±0.79 while that in vehicle group was 5.6±0.40, indicating that FI-RSV significantly enhanced lung pathological damage (Fig. 6e and Fig. 6f). This result demonstrated that while FI-RSV could provide protection again RSV infection, it elicited significant vaccine related disease, consistent with previous report.[24, 25]

Compared with the vehicle group and FI-RSV group, STR-V003 induced dose-dependent RSV preF-specific IgG antibodies (Fig. 6b) and significantly higher levels of neutralizing antibodies at all dose levels (Fig. 6c). Compared with the vehicle group, STR-V003 significantly reduced the RSV viral loads by 2.75, 2.67, and 2.58 Log10 (Copies/g lung) in the lung tissue of animals treated with 1.25, 5, or 20 μg vaccine, respectively (Fig. 6d). As shown in Fig. 6e and Fig. 6f, test vaccine STR-V003 (1.25 µg/mouse) significantly reduce perivasculitis, peribronchiolitis and alveolitis inflammation scores, and a medium dose (5 µg/mouse) significantly reduce alveolitis inflammation. High dose (20 µg/mouse) significantly reduce peribronchial and alveolitis inflammation scores. The experimental results showed that the tested vaccine had no enhancement of lung damage. Taken together, these results demonstrated that STR-V003 provided efficient protection again RSV infection without causing VED.

### Immunogenicity and challenge protection of STR-V003 in cotton rat model

The immunogenicity and challenge protection of STR-V003 were further evaluated in cotton rat model. Cotton rats were immunized with STR-V003 vaccine (containing 5 or 20 μg mRNA, respectively) via IM injections on Day 0 and Day 21. On Day 50, RSV/A2 (ATCC, VR-1540) or RSV/B9320 (ATCC, VR-955) were inoculated intranasally (1.5×10^6^ PFU/animal) (Supplementary Table 3). Serum samples were collected on Day 49 for neutralizing antibody. The turbinate bones and lung tissues were collected on Day 54 after the animals were euthanized (Fig. 7a).

**Figure 7.**
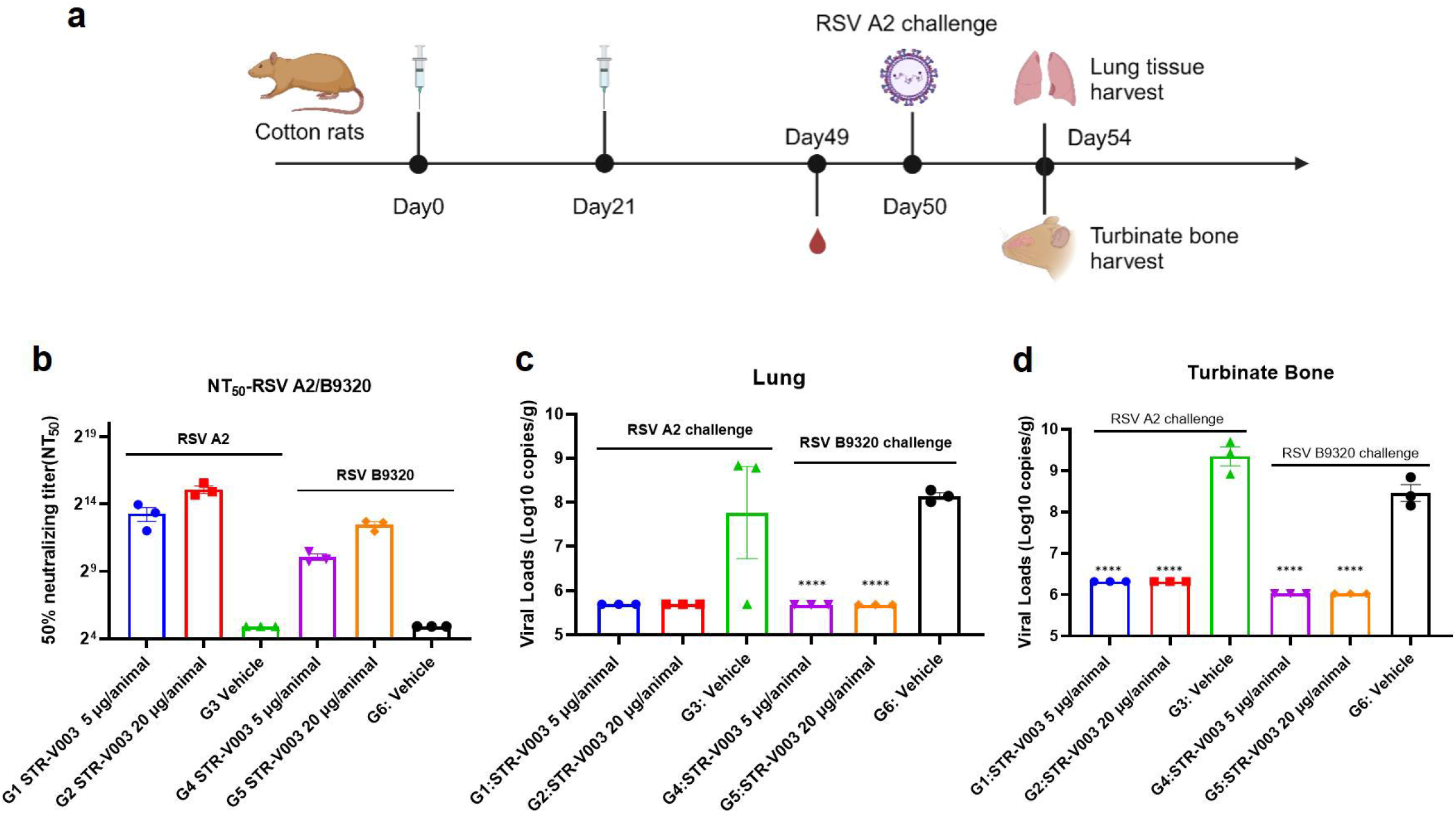
Protection Against RSV Challenge in Cotton Rat Model. **(a)** The schematic diagram of immunogenicity and challenge protection of STR-V003 in cotton rat model. Cotton rats were immunized with STR-V003 vaccine (containing 5 or 20 μg mRNA, respectively) via IM injections on Day 0 and Day 21. On Day 30, RSV/A2 (ATCC, VR-1540) or RSV/B9320 (ATCC, VR-955) were inoculated intranasally (1.5×10^6^ PFU/animal). Serum samples were collected on Day 49. The turbinate bones and lung tissues were collected on Day 54 after the animals were euthanized. **(b)** The 50% neutralizing titer (NT50) of RSV A2 and RSV B9320 in cotton rats on Day 49. (**c, d)** Viral loads in lung tissue and turbinate bone on Day 54. **(e)** H&E staining of lung tissues in cotton rats on Day 54.

Neutralizing antibody was measured by microneutralization to evaluate the immunogenicity of STR-V003. To evaluate the efficacy of the vaccine in vivo, RSV viral loads in lung tissues and turbinate bones were measured by RT-qPCR. Compared with the vehicle, the tested vaccine STR-V003 induced dose-dependent levels of neutralizing antibodies (against RSV/A2 or RSV/B9320) (Fig. 7b), significantly reduced the RSV viral loads in the lung tissue (by 2.081 Log10 (copies/g lung) for RSV A2, by 2.462 Log10 (copies/g lung) for RSV B9320 at both doses) and turbinate bone (by 3.023 Log10 (copies/g bone) for RSV A2, by 2.433 Log10 (copies/g bone) for RSV B9320 at both doses) of infected animals (Fig. 7c and Fig. 7d). The experimental results showed that test vaccine STR-V003 had good anti-RSV virus effect in vivo.

Evaluation of lung tissue lesions by HE staining, perivascular, peribronchial, alveolar and pulmonary interstitial inflammation scores. The results are shown in Fig. 7e and Supplementary Table4. Compared with the vehicle group (G3), test vaccine STR-V003 group (G1-2) had no enhancement of lung damage. Compared with the vehicle group (G6), test vaccine STR-V003 group (G3-4) had no enhancement of lung damage. The experimental results showed that the test vaccine STR-V003 had no enhancement of lung damage, demonstrating efficient protection against RSV infection was provided by STR-V003 without causing VED.

### Safety of STR-V003 in SD rat

Safety studies on STR-V003 and A1-EP10-O18A (as empty LNPs) were designed in accordance with the WHO guidelines on vaccines.[26] As shown in Fig 8a, the potential sub-chronic toxic effects and the reversibility, persistence, or any delayed occurrence of observed adverse events were evaluated for STR-V003 and empty LNPs in a 4-week repeat-dose toxicology study with a 2-week recovery phase in SD rats. The rats were randomly assigned into 7 groups (10/sex/group) and received negative control (Sodium Chloride Injection), STR-V003 at 1, 1.5, or 2 doses (50, 75, or 100 µg), or empty LNPs at 0.5, 0.75, or 1 mL (equivalent to 1, 1.5, or 2 vaccine doses), via IM injections (Q2W, for a total of 3 administrations). On Day 32, half of the animals (5/sex/group) were euthanized and necropsied while the remaining animals (5/sex/group) were euthanized on Day 43. Blood samples for hematology determinations were collected on Day 4, on Day 32 and Day 43.

**Figure 8.**
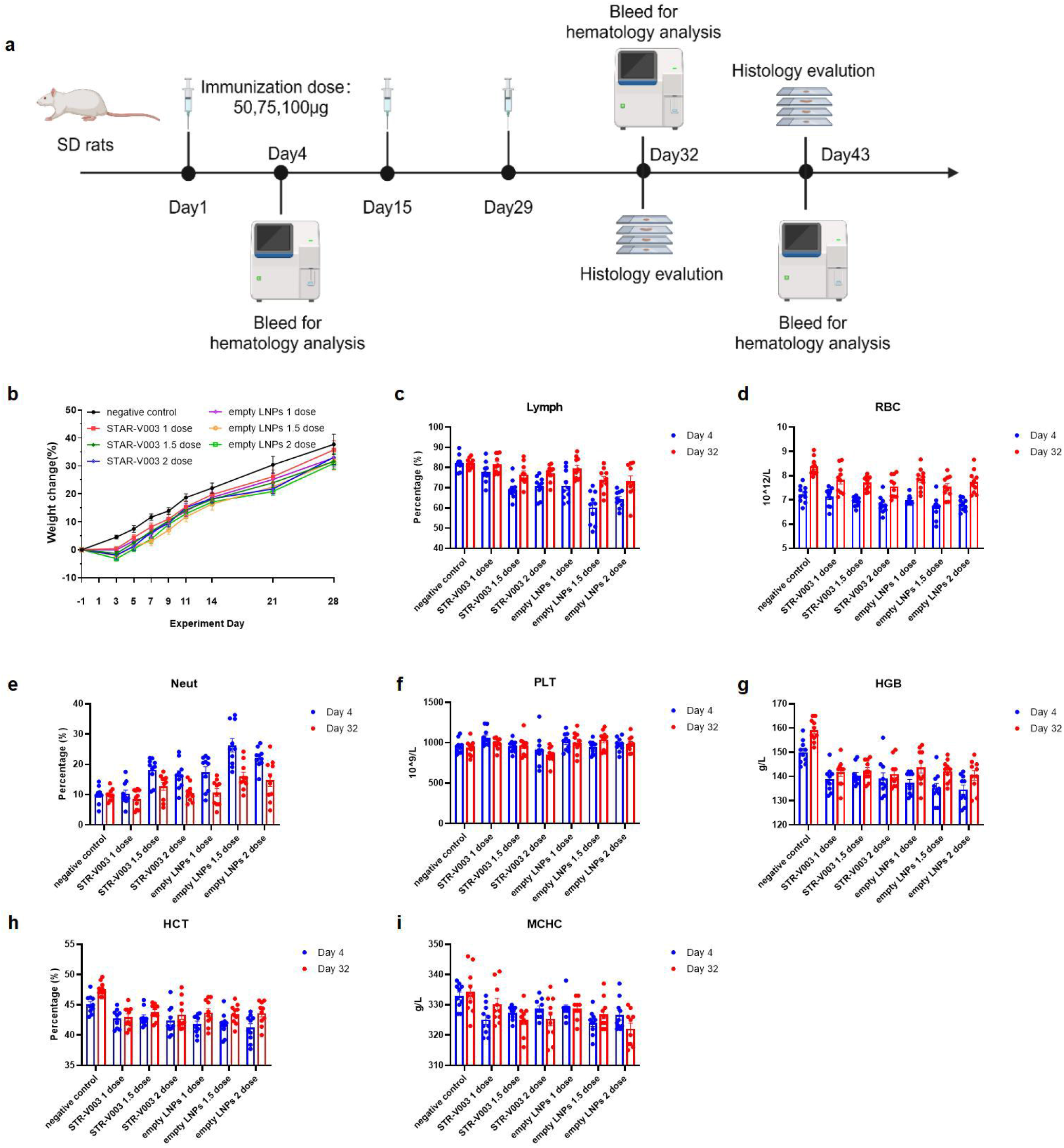
Safety of STR-V003 in SD rat. **(a)** The schematic diagram of safety studies on STR-V003 and A1-EP10-O18A in SD rats. The SD rats received negative control (Sodium Chloride Injection), STR-V003 at 1, 1.5, or 2 doses (50, 75, or 100 µg), or empty LNPs at 0.5, 0.75, or 1 mL (equivalent to 1, 1.5, or 2 vaccine doses), via IM injections (Q2W, fr a total of 3 administrations) following a 2-week recovery period. **(b)** Body weight changes of rats after immunization. **(c-i)** Pathological indicators in the blood of rats at Day4 and Day32, including lymphocyte (Lymph) percentage (**c**), Erythrocyte count (RBC) (**d**), Neutrophilic granulocyte (Neut) percentage (**e**), platelet (PLT) counts (**f**), hemoglobin (HGB) concentration (**g**), hematocrit (HCT) percentage (**h**), and mean corpuscular hemoglobin concentration (MCHC) (**i**).

All doses of STR-V003 and empty LNPs were tolerated. There was no treatment-related mortality or moribundity. Local irritation reactions were observed at the injection sites, with a trend of recovery after a 2-week recovery period.

All doses of STR-V003 and empty LNPs were tolerated. There was no treatment-related mortality or moribundity. No empty LNPs / STR-V003 related changes in ophthalmoscopic examinations throughout the study. Swelling in the injection sites was noted in animals in all empty LNPs groups and the STR-V003 groups on 2 to 4 days after dosing, which recovered afterwards.

Compared with the negative control group of the same gender in the same period, decreased body weight and body weight gain were observed at 1 week after each dosing in the STR-V003 and empty LNPs groups (Fig 8b). A trend toward recovery was seen at Week 2 after dosing. These lower body weights were considered secondary changes to local irritancy of the dose. At the same dose, the change range were generally similar in the empty LNPs and the STR-V003 groups.

Compared with the negative control group of the same gender in the same period, increased Neut, and decreased RBC, HGB and HCT in rats were observed on Day 4 and Day 32 in the STR-V003 and empty LNPs groups at 1,1.5 and 2 dose/animal (Fig 8c-Fig 8i). These changes were considered possibly related to an immune response and/or local irritation and/or the acute phase response.[27] A trend towards recovery or recovery was seen after 2 weeks of drug withdrawal (Day 43). At the same dose, the change range were similar in the empty LNPs and the STR-V003 groups.

The animals were euthanized at scheduled necropsy for macroscopic changes, organ weight and histopathological examination of liver, submandibular lymph nodes, mesenteric lymph nodes, inguinal lymph nodes, spleen and thymus. At 3 days after the last dosing (Day 32), decreased the organ weight, organ-to-body/ brain weight ratios of the thymus and increased the organ weight, organ-to-body/ brain weight ratios of the spleen were observed in the STR-V003 and empty LNPs groups at 1, 1.5 and 2 dose/animal (Supplementary Table 5). As shown in Table 1, microscopically, minimal to moderate cellularity lymphocytes decreased of cortex/medulla in the thymus and minimal to moderate vacuolation of hepatocytes around portal areas in the liver were observed in the STR-V003 and empty LNPs groups, minimal to slight cellularity lymphocytes increased in the white pulp in the spleen was observed in the STR-V003 and empty LNPs groups at 1.5 and 2 dose/animal, extramedullary hematopoiesis increased of the red pulp in the spleen was observed in the STR-V003 groups at 1, 1.5 and 2 dose/animal and empty LNPs groups at 1.5 and 2 dose/animal. A trend towards recovery or recovery was seen after 2 weeks of drug withdrawal (Day 43). At the same dose, the change range, incidence and severity of lesions were similar in the empty LNPs and the STR-V003 groups.

**Table 1.**
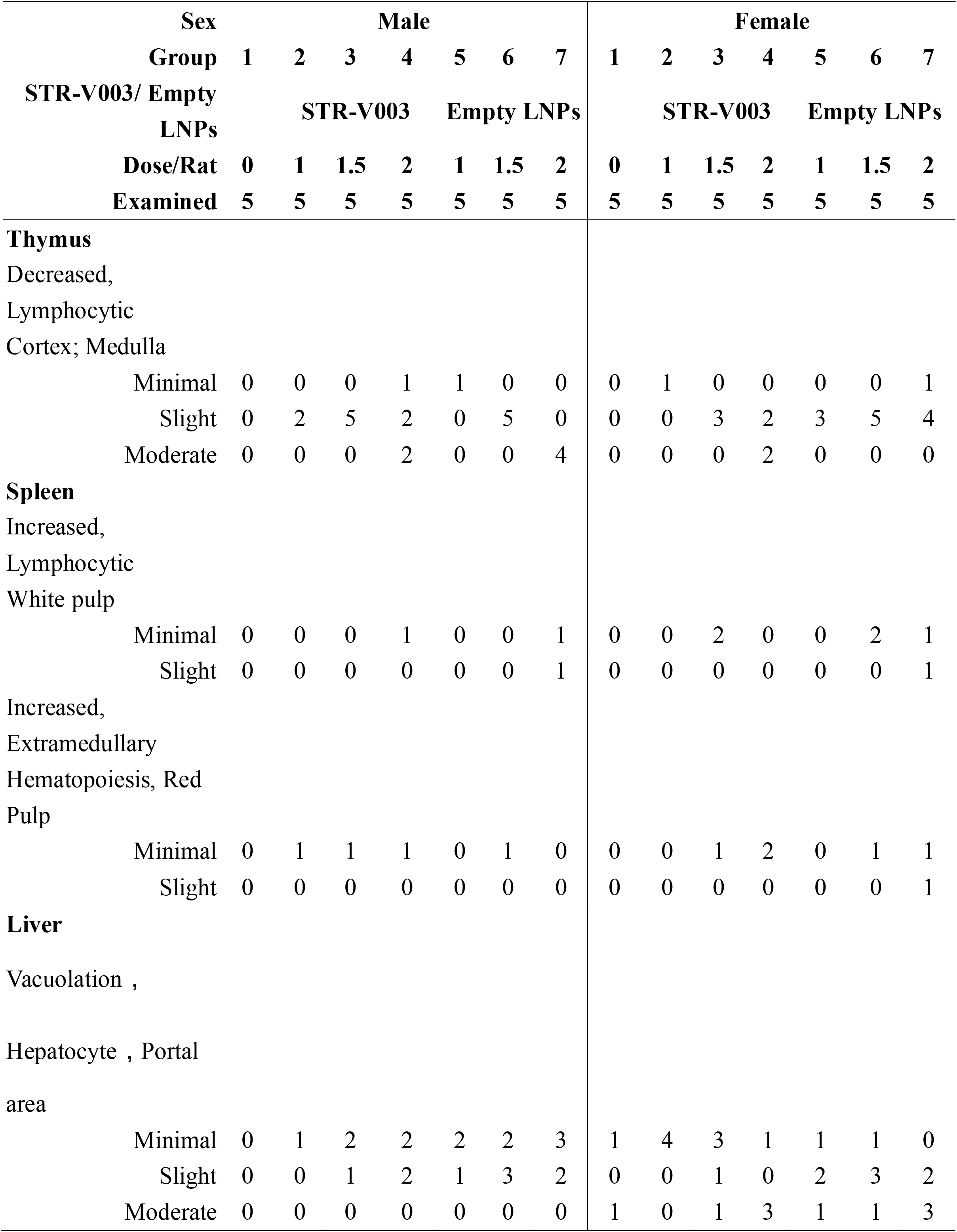
Empty LNPs / STR-V003-related microscopic findings, Terminal necropsy (Day 32)s.

## Discussion

In this study, STR-V003 is an mRNA vaccine produced by encapsulating the mRNA that encodes RSV preF protein in lipid nanoparticles (LNPs). A series of studies has been conducted to assess the immunogenicity and challenge protection of of STR-V003.

The in vitro and in vivo pharmacology studies screened the mRNA sequences encoding RSV preF protein based on protein expression levels in transfected cell lines and immunogenicity in mice. The mRNA sequence (V003-210-T-plus) was determined to be the optimal sequence to be used in STR-V003.

Currently, there are no animal models that can fully recapitulate the pathogenesis of RSV infection in humans. The non-human primates (except chimpanzees) are only semi-permissive for hRSV replication and exhibit little or no clinical signs of disease when infected with hRSV. Even though cotton rats and mice are also only semi-permissive for hRSV, these two models have been informative in providing insights of RSV pathogenesis and mechanisms of immunity. These two models were thus selected to study RSV challenge protection by STR-V003. STR-V003 administered via IM injections on Day 1 and Day 21 prior to intranasal inoculation of RSV/A2 (in both models) or RSV/B9320 (in cotton rats) led to dose-dependent induction of RSV-neutralizing antibody (lasted for at least 7 weeks in cotton rats and 5 weeks in mice after 1^st^ vaccination) and RSV preF-binding antibody (lasted for at least 4 weeks after 1^st^ vaccination). A significant reduction in tissue viral loads was observed in both models. Both the immune responses and the protection induced by STR-V003 were stronger than those induced by FI-RSV. Importantly, unlike FI-RSV, STR-V003 did not cause enhanced lung pathology. Taken together, these results demonstrated that STR-V003 is able to induce robust and lasting immune responses and provide significant protection against RSV infection without inducing vaccine enhanced disease (VED). It is noteworthy that STR-V003 vaccination induced both RSV A- and RSV B-neutralizing antibodies and reduced the viral loads of both RSV A and RSV B strains, suggesting that STR-V003 is capable to protect infection of both RSV A and RSV B strains, the two major groups of RSV that co-circulate worldwide[28].

The safety studies did not identify unexpected toxicities following repeated IM injections of STR-V003 ((50, 75, or 100 µg, Q2W, 3 administrations in total) in SD rats. The changes observed were considered to be the expected immune responses to mRNA vaccine or secondary to inflammation. The local irritation at the injection sites observed in SD rats in safety studies was reversible. As toxicology studies indicated that STR-V003 did not affect vital physiological functions (e.g., central nervous system, respiratory, and cardiovascular functions) other than the immune system.

In conclusion, a comprehensive panel of studies were conducted to evaluate the immunogenicity, protection efficacy and safety profiles of STR-V003. The above results showed that the test vaccine STR-V003 had good immunogenicity and in vivo antiviral efficacy, had no disease enhancement effect. The scope of the toxicological evaluation of STR-V003 demonstrated an acceptable safety profile. The data obtained from these studies support the further human trials of STR-V003.

## Materials & methods

### Cells and viruses

293T cells were purchased from Cell Bank/Stem Cell Bank, Chinese Academy of Sciences, HT1080 cells were purchased from Nanjing Cobioer Biosciences, A549 cells were donated from Westlake University. 293T, HT1080 and A549 cells were grown in complete medium (DMEM medium supplemented with 10% FBS) at 37□ in 5% CO_2_. FI-RSV is produced by WuXi Discovery Biology. Respiratory syncytial virus, RSV/A2 (ATCC, VR-1540) and RSV/B9320 (ATCC, VR-955) were used for detecting of neutralizing antibody.

### Preparation of mRNA

In vitro transcription was performed using the TranscriptAid T7 High Yield Transcription Kit (Thermo K0441), using linear DNA as a template for generating mRNA, with addition of a certain proportion of pseudouridine and capping reagents. In vitro transcription conditions: the reaction system was configured according to the kit instructions, reaction at 37 °C for 0.5-2 hours, the transcript was digested for 30 minutes by DNase, and the transcript was purified with Monarch RNA Cleanup Kit (NEB T2040L).

### Lipid synthesis

All lipids are synthesized by a similar method.

#### 1-((3-((3-aminopropyl)(methyl)amino)propyl)amino)decan-2-ol

2-octyloxirane (12.0 g, 1.0 eq.) was added to a solution of N1-(3-aminopropyl)-N1-methylpropane-1,3-diamine (44.62 g, 4.0 eq.) in EtOH (446.2 mL, 10V), then the mixture was heated to 70 oC and stirred for 15h. TLC showed the reaction was completed. The solution was evaporated under reduced pressure. H2O (500 mL) was then added to the residue, and the solution was washed by DCM (500 mL x 5), then the combined organic phase was washed by saturated brine (300 mL) and dry over Na2SO4, after removing the solvent under vacuum, compound 3 was obtained as a colorless oil (18 g, 77.74% yield). 1H NMR (400 MHz, CDCl3) δ 3.60 (dddd, J = 9.8, 7.2, 4.4, 2.9 Hz, 1H), 2.79 – 2.57 (m, 5H), 2.47 – 2.34 (m, 5H), 2.20 (s, 3H), 1.64 (dp, J = 18.7, 6.9 Hz, 4H), 1.49 – 1.18 (m, 14H), 0.93 – 0.81 (m, 3H).

#### (9Z,12Z)-octadeca-9,12-dien-1-yl acrylate

acryloyl chloride (9.76 g, 1.15 eq.) was added to a solution of (9Z,12Z)-octadeca-9,12-dien-1-ol (25.0 g, 1.0 eq.) in DCM (1250 mL, 50 V) at 0 °C, then the mixture was stirred at 0 °C for 0.5 h. A solution of Et_3_N (14.24 g, 1.5 eq.) in DCM (50 mL) was added and stirred at 0 °C for 2 h. TLC showed the reaction was completed. H_2_O (500 mL) was then added to the mixture, the organic phase was separated and the aqueous phase was extracted by DCM (500 mL x 3), then the combined organic phase was washed by saturated brine (300 mL) and dry over Na_2_SO_4_, after removing the solvent under vacuum, the crude product was purified by silica-gel chromatography (DCM/hexane = 1:100, v/v) to give compound 6 as a colorless oil (29 g, 96.44%). 1H NMR (400 MHz, CDCl_3_) δ 6.40 (dd, J = 17.3, 1.5 Hz, 1H), 6.12 (dd, J = 17.4, 10.4 Hz, 1H), 5.81 (dd, J = 10.4, 1.5 Hz, 1H), 5.49 – 5.26 (m, 4H), 4.15 (t, J = 6.7 Hz, 2H), 2.77 (t, J = 6.4 Hz, 2H), 2.05 (q, J = 6.8 Hz, 4H), 1.72 – 1.62 (m, 2H), 1.41 – 1.24 (m, 17H), 0.89 (t, J = 6.8 Hz, 3H).

#### di((9Z,12Z)-octadeca-9,12-dien-1-yl)

##### 3,3’-((3-((3-((2-hydroxydecyl)(3-(((9Z,12Z)-octadeca-9,12-dien-1-yl)oxy)-3-oxopropyl)amino) propyl)(methyl)amino)propyl)azanediyl)dipropionate

a solution of (9Z,12Z)-octadeca-9,12-dien-1-yl acrylate (18.6 g, 3.5 eq.) and 1-((3-((3-aminopropyl)(methyl)amino)propyl)amino)decan-2-ol (5.0 g, 1.0 eq.) was stirred at 20 °C for 1h under N_2_, then BHT (365.41 mg, 0.1 eq.) and AcOH (49.79 mg, 0.05 eq.) was added and stirred at 70 °C for 80 h. TLC showed the reaction was completed. The mixture was purified by silica-gel chromatography (DCM/MeOH = 30:1, v/v) to give compound A as a yellow oil (43g, yield 57.03%). ^1^H NMR (400 MHz, CDCl_3_) δ 5.45 – 5.24 (m, 12H), 4.11 – 3.98 (m, 6H), 3.63 – 3.53 (m, 1H), 2.95 – 2.84 (m, 1H), 2.81 – 2.53 (m, 12H), 2.49 – 2.15 (m, 18H), 1.67 – 1.53 (m, 10H), 1.51 – 1.15 (m, 65H), 0.88 (td, *J* = 6.8, 4.8 Hz, 12H). HR-MS: [M+H]^+^= 1263.15105

### Lipid nanoparticle preparation

mRNA was dissolved in citrate buffer (pH 4) and adjusted the concentration of mRNA to 0.2 mg/mL, so that obtaining aqueous layer thereby. A1-EP10-O18A (synthesized according to our patented method), 1,2-distearoyl-sn-glycero-3-phosphocholine (DSPC), cholesterol and DMG-PEG2000 were dissolved in desired molar fractions in dry ethanol and the total concentration of lipids was adjusted to 10 mg/mL, thereby obtaining organic layer. The aqueous layer and organic layer were admixed in 3:1 ratio (v/v) by microfluidic device (NanoAssemblr® Ignite™) at total flow rate of 12 mL/min. The mixture was 10-fold diluted with PBS buffer (pH 7.4). Ethanol was separated by tangential flow filtration (Repligen, TFF). The solution was concentrated to 0.1 mg/mL (mRNA concentration) and filtrated by 0.22 μm millipore filter to afford mRNA containing lipid nanoparticles.

### mRNA transfection

293T, HT1080 and A549 cells were inoculated in 6-well or 24-well plates. Twenty-four hours later, the cells were transfected with mRNA of RSV preF protein using Lipofectamine 2000 (Invitrogen). Cells were collected 24 hours after transfection. The expression of RSV preF protein was then detected by western blot with a monoclonal antibody that recognizes RSV F protein (Sinobiological,11049-R302), FACS with F488 Anti-Respiratory Syncytial Virus Fusion Protein Antibody(Φ) (recognize the antigenic site Φ of RSV preF protein, Vazyme), RSV F Antibody (4D5) (Epitope I, novoprotein, DA102), RSV F Antibody (11A9) (Epitope II, novoprotein, DA091), RSV F Antibody (4B9) (Epitope III, novoprotein, DA110), RSV F Antibody (9F4) (Epitope IV, novoprotein, DA118), RSV Pre-F Antibody (7E11) (Epitope V, novoprotein, DA119), RSV Pre-F Antibody (3C12) (Epitope Φ, novoprotein, DA101) and Rabbit Anti-Human IgG H&L (FITC) (abcam, ab6755), and ELISA with Respiratory Syncytial Virus preF Antibody Pair (Φ & □) (Vazyme).

### Animal studies

In the study of *in vivo* screening of RSV preF mRNA sequence for STR-V003, BALB/c mice (6∼8weeks old) in each group were injected with 50 µL mRNA-LNPs twice on Day0 and Day14. Vehicle was used as negative control. Body weight was measured every 4 days until the end of the study. Serum samples were collected on days 14 and 35 for detection of RSV preF-binding IgG antibody (Vazyme, DD3910-01).

In the study of immunogenicity and challenge protection of STR-V003 in mouse model, female and male, 6-8 weeks old, BALB/c mice were immunized with vehicle, test vaccine STR-V003, FI-RSV by IM route on day 0 and day 21. Mice were deeply anesthetized by intraperitoneal injection of anesthetic (30 mg/kg Zoletil 50 and 6 mg/kg Xylazine Hydrochloride) on the day of inoculation (day 35), and then inoculated with the RSV/A2 via intranasal inhalation. The inoculation amount is 1.0E+06 PFU/50 μL/mouse. Serum samples were collected on days 14, 28, and day 35 for detection of RSV preF-specific IgG antibody and Neutralization antibody titers. On the 40th day, the animals were euthanized by overdose anesthesia and exsanguination, the lung tissue samples were collected for detection of viral loads and lesions of lung tissue. After immunization, mice were weighed once a day for 6 consecutive days until the abnormal symptoms have significantly improved and recovered, and then three times a week. After virus infection, mice were weighed once a day to observe and record their health and survival status.

In the study of immunogenicity and challenge protection of STR-V003 in cotton rat model, 5-7 weeks old, cotton rat were immunized with vehicle, test STR-V003 by IM route on day 0 and day 21, and anesthetized with isoflurane gas in a gas anesthesia machine on the day of inoculation (day 50). Each animal was intranasally instilled with 50 μL of viral solution, and the animals were observed to inhale the viral solution, return to the home cage after waking up, and closely observe the status of the animals. The inoculation amount is 1.5E+06 PFU/50 μL/cotton rats. Serum samples were collected on days 49 for neutralizing antibody. The turbinate bones and lung tissues were collected on day 54 after the animals were euthanized for detection of viral loads and lesions of lung tissue. Animals were weighed and recorded three times a week before infection and once a day after infection.

The protocol and any amendments or procedures involving the care and use of all animals in this study will be reviewed and approved by the Institutional Animal Care and Use Committee (IACUC) of before conducting the study. During the study, the care and use of animals will be conducted in accordance with the regulations of the Association for Assessment and Accreditation of Laboratory Animal Care (AAALAC).

### Evaluation of RSV preF/F-specific antibody titers

In the study of in vivo screening of RSV preF mRNA sequence for STR-V003, the titers of serum specific IgG antibody against RSV preF of mouse were tested using an ELISA kit (Vazyme, DD3910-01) according to the manufacturer’s instructions.

Evaluation of RSV F-specific antibody titers were conducted as previously described.[29] ELISA plates were coated with recombinant human RSV fusion glycoprotein (total-F protein, final concentration: 0.2μg /mL) (Sino Biological Inc. 11049-V0813). The coated plates were incubated with a given serum dilution, and anti-mouse antibodies labeled with HRP (Abcam, ab6728.) were used to measure the binding of antibodies specifically to RSV-F protein using TMB substrate.

In mouse model challenge experiment, the titers of serum specific IgG antibody against RSV preF of mouse infection experiments were measured by ELISA. Briefly, HRSV(A) prefusion glycoprotein F0 (Acro Biosystems, RSF-V52H7) was coated on a microplate prior to addition of titrated serum. Wells were washed and detected with goat anti-mouse IgG-HRP (Abcam, ab97023) and TMB substrate. The absorbance values at 450 nm were measured using a multi-functional microplate reader, and the RSV preF IgG antibody titers of serum samples of each animal were analyzed using a four-parameter fitting method.

### Neutralizing antibody

RSV neutralizing antibody was measured to evaluate the immunogenicity of STR-V003. Neutralizing antibody was detected by microneutralization. Briefly, serial dilutions of heat-inactivated serum from vaccinated or control animals were incubated with virus for 2 hours at 37°C. The serum-virus mixture or virus alone was added to the wells containing Hep-2 cells and incubated for 2h or 5 days at 37°C. After removing the supernatant, culture medium was added and culture was continued for about 22 hours. The cells were then fixed and virus-specific foci were detected by antibody against RSV F protein (Sino Biological, 11049-R338) or RSV G protein (Vazyme, DD1606), and secondary antibody (HRP-labeled Goat Anti-Rabbit IgG (H+L). The signal was developed by TMB and absorbance was read at 450 nm. NT50 (The reciprocal of the serum dilution ratio when RSV viruses suppressed by 50%) was calculated to indicate the neutralizing activity of antibody against RSV viruses.

### Quantification of viral RNA in the lung tissue or turbinate bones of the infected animal by RT-qPCR

To evaluate the efficacy of the vaccine in mouse model challenge experiment, RSV viral loads in lung tissue were measured by RT-qPCR. Briefly, RNA was extracted from the tissue lysates using RNeasy Mini Kit. The cDNA was synthesized using a commercial kit (Thermo Fisher Scientific, 4368814) and the RSV-specific reverse transcription primers. The qPCR was performed using FastStart Universal Probe Master (Roche, 04914058001). The RSV copy numbers of samples were calculated based on a standard curve made using plasmid RSV-A-N copy numbers and qPCR amplification cycle numbers.

In cotton rats model challenge experiment, RSV viral loads in lung tissues and turbinate bones were measured by RT-qPCR. Briefly, RNA was extracted from ground tissues using DNA/RNA Extraction Kit (Vazyme, RM201-02). cDNA synthesis and PCR were performed in a single tube using HiScript II U+ One Step qRT-PCR Probe Kit (Vazyme, Q223-01). The RSV copy numbers of samples were calculated based on a standard curve made using plasmid RSV-A-N copy number and qPCR amplification cycle number.

### Histopathology assay

The left lung was perfused with 0.2 mL of 4% paraformaldehyde and fixed in 4% paraformaldehyde. Fixed samples were embedded in paraffin, sectioned, and stained with HE. Finally, lung sections should be scored by a person blinded to the group assignment.

### Safety Study of STR-V003/Empty LNPs

A total of 140 SD rats (70 animals/sex) were randomly assigned to 14 groups with 5 animals/sex/group in Groups 1 to 7 designated as the main study groups, and 5 animals/sex/group in Groups 8 to 14 as the satellite groups. Rats were treated with Sodium Chloride Injection (1 mL/animal) as negative control for Group 1 and 8, STR-V003 (mRNA content of 50 μg/dose) at doses of 0.5 mL/1, 0.75 mL/1.5 and 1 mL/2 dose/animal for animals of Groups 2 and 9, Groups 3 and 10 or Groups 4 and 11, respectively, empty LNPs at doses of 0.5 mL/1, 0.75 mL/1.5 and 1 mL/2 dose/animal for animals of Groups 5 and 12, Groups 6 and 13 or Groups 7 and 14, respectively. The animals were administered via intramuscular injection once every two weeks for 4 consecutive weeks (3 doses in total).

Parameters evaluated in this study included clinical observations (including administration site observations), body weight, food consumption, body temperature, hematology. The animals in Groups 1 to 7 and Groups 8 to 14 were euthanized 3 days after the last dosing (Day 32) and after the end of recovery period (Day 43) for macroscopic changes, organ weight and histopathological examination of liver, submandibular lymph nodes, mesenteric lymph nodes, inguinal lymph nodes, spleen, thymus.

### Statistical analysis

GraphPad Prism 7.00 was used for statistical analysis of the data. One-way ANOVA was used to analyze the statistical difference of lung tissue viral loads. One-way ANOVA and Two-way ANOVA were used to analyze the statistical difference of lung tissue pathology.

### Safety Study of STR-V003/Empty LNPs

All statistical tests were conducted as 2-sided tests, and the level of significance were set at 5% or P≤0.05. Group means and standard deviations (SD) were calculated for males and females, respectively, in the Provantis system (SAS 9.4 software), including body weights, body weight gain, food consumption (except the recovery period), body temperature, organ weights, organ-to-body/brain weight ratios, clinical pathology (hematology, coagulation, clinical chemistry), bone marrow smears, cytokines, phenotype of immune cells, and bone marrow smears, etc. in the negative control article and dose groups. The data were analyzed with the following procedures: A Levene’s test were performed on the original data to test for variance homogeneity. When the result shows no significance (P>0.05), a one-way analysis of variance (ANOVA) were performed. If ANOVA shows significance (P≤0.05), Dunnett’s test (parametric method) were performed for multiple comparisons.

In the case of variance heterogeneity (P≤0.05 by Levene’s test), the data were logarithmically transformed using the natural logarithm (ln transformation), then the transformed data were tested for variance homogeneity as described above using Levene’s test. In the case of variance heterogeneity of transformed data (P≤0.05 by Levene’s test) or contains negative values, a Kruskal-Wallis test (non-parametric method) were performed on the original data. If Kruskal-Wallis test shows significance (P≤0.05), a two-independent-sample test (Mann-Whitney U Wilcoxon) were performed for multiple comparisons.

When a dataset has zero values, the zero values were regarded as 1/10 of the smallest positive value in the dataset when logarithmical transformation is performed.

## Acknowledgements

We thank employees of Starna Therapeutics for their helpful technical and scientific support with animal study and materials preparation. We would like to thank Dr Qiang Cheng for kindly providing suggestions and advice. This study was supported in part from Starna Therapeutics (STR-V003), Jiangsu Provincial Science and Technology Project (SBK2023070025), Suzhou Science and Technology Project (ZXL2023267). National Center of Technology Innovation for Biopharmaceuticals (NCTIB2023XB02001).

## Author contribution

R.H., H.B., Q.L., X.Y., conceived the project and designed the experiments and wrote the manuscript. R.H. and X.Y. supervised the project. Q.L. carried out the mRNA synthesis. Y. Z. prepared the LNP and quality control. Y.G and Y.Z. performed the molecular experiments. All the authors discussed the results and commentated manuscript.

## References

1. Adhikari, B., et al., A multi-center study to determine genetic variations in the fusion gene of respiratory syncytial virus (RSV) from children <2 years of age in the U.S. J Clin Virol, 2022. 154: p. 105223.

2. Kim, S., et al., RSV genomic diversity and the development of a globally effective RSV intervention. Vaccine, 2021. 39(21): p. 2811–2820.

3. Battles, M.B. and J.S. McLellan, Respiratory syncytial virus entry and how to block it. Nature Reviews Microbiology, 2019. 17(4): p. 233–245.

4. Andreano, E., et al., The respiratory syncytial virus (RSV) prefusion F-protein functional antibody repertoire in adult healthy donors. EMBO Mol Med, 2021. 13(6): p. e14035.

5. Espeseth, A.S., et al., Modified mRNA/lipid nanoparticle-based vaccines expressing respiratory syncytial virus F protein variants are immunogenic and protective in rodent models of RSV infection. NPJ Vaccines, 2020. 5(1): p. 16.

6. Mejias, A., et al., Respiratory Syncytial Virus Vaccines: Are We Making Progress? The Pediatric infectious disease journal, 2019. 38(10): p. e266–e269.

7. Domachowske, J.B., E.J. Anderson, and M. Goldstein, The Future of Respiratory Syncytial Virus Disease Prevention and Treatment. Infect Dis Ther, 2021. 10(Suppl 1): p. 47–60.

8. Taleb, S.A., et al., Human respiratory syncytial virus: pathogenesis, immune responses, and current vaccine approaches. Eur J Clin Microbiol Infect Dis, 2018. 37(10): p. 1817–1827.

9. Ruckwardt, T.J., K.M. Morabito, and B.S. Graham, Immunological Lessons from Respiratory Syncytial Virus Vaccine Development. Immunity, 2019. 51(3): p. 429–442.

10. Topalidou, X., A.M. Kalergis, and G. Papazisis, Respiratory Syncytial Virus Vaccines: A Review of the Candidates and the Approved Vaccines. Pathogens, 2023. 12(10).

11. Syed, Y.Y., Respiratory Syncytial Virus Prefusion F Subunit Vaccine: First Approval of a Maternal Vaccine to Protect Infants. Paediatr Drugs, 2023. 25(6): p. 729–734.

12. Bonneux, B., et al., Direct-acting antivirals for RSV treatment, a review. Antiviral Research, 2024. 229: p. 105948.

13. McLellan, J.S., et al., Structure-based design of a fusion glycoprotein vaccine for respiratory syncytial virus. Science, 2013. 342(6158): p. 592–8.

14. Hsieh, C.-L., et al., Structure-based design of prefusion-stabilized SARS-CoV-2 spikes. Science, 2020. 369(6510): p. 1501–1505.

15. Krarup, A., et al., A highly stable prefusion RSV F vaccine derived from structural analysis of the fusion mechanism. Nat Commun, 2015. 6: p. 8143.

16. Bolt, G., L.O. Pedersen, and H.H. Birkeslund, Cleavage of the respiratory syncytial virus fusion protein is required for its surface expression: role of furin. Virus Res, 2000. 68(1): p. 25–33.

17. Smith, G., et al., Respiratory syncytial virus fusion glycoprotein expressed in insect cells form protein nanoparticles that induce protective immunity in cotton rats. PLoS One, 2012. 7(11): p. e50852.

18. Earp, L.J., et al., The many mechanisms of viral membrane fusion proteins. Curr Top Microbiol Immunol, 2005. 285: p. 25–66.

19. Zimmer, G., L. Budz, and G. Herrler, Proteolytic activation of respiratory syncytial virus fusion protein. Cleavage at two furin consensus sequences. J Biol Chem, 2001. 276(34): p. 31642–50.

20. González-Reyes, L., et al., Cleavage of the human respiratory syncytial virus fusion protein at two distinct sites is required for activation of membrane fusion. Proc Natl Acad Sci U S A, 2001. 98(17): p. 9859–64.

21. Park, J.H. and H.K. Lee, Delivery Routes for COVID-19 Vaccines. Vaccines (Basel), 2021. 9(5).

22. Byrd, L.G. and G.A. Prince, Animal models of respiratory syncytial virus infection. Clin Infect Dis, 1997. 25(6): p. 1363–8.

23. Taylor, G., Animal models of respiratory syncytial virus infection. Vaccine, 2017. 35(3): p. 469–480.

24. Boelen, A., et al., Both immunisation with a formalin-inactivated respiratory syncytial virus (RSV) vaccine and a mock antigen vaccine induce severe lung pathology and a Th2 cytokine profile in RSV-challenged mice. Vaccine, 2000. 19(7-8): p. 982–91.

25. Waris, M.E., et al., Respiratory synctial virus infection in BALB/c mice previously immunized with formalin-inactivated virus induces enhanced pulmonary inflammatory response with a predominant Th2-like cytokine pattern. Journal of Virology, 1996. 70(5): p. 2852–2860.

26. Organization, W.H. Annex 1 WHO guidelines on nonclinical evaluation of vaccines. 2005; Available from: https://www.who.int/publications/m/item/nonclinical-evaluation-of-vaccines-annex-1-trs-no-927.

27. Agency, E.M. Assessment report: Comirnaty. 2021; Available from: https://www.ema.europa.eu/en/documents/assessment-report/comirnaty-epar-public-assessment-report_en.pdf.

28. Muñoz-Escalante, J.C., et al., Respiratory syncytial virus B sequence analysis reveals a novel early genotype. Scientific Reports, 2021. 11(1): p. 3452.

29. Zhang, L., et al., Design and characterization of a fusion glycoprotein vaccine for Respiratory Syncytial Virus with improved stability. Vaccine, 2018. 36(52): p. 8119–8130.

