## supporting data for "Efficacy and safety of an mRNA-based RSV preF vaccine in preclinical studies"

**Supplementary information**

**Supplementary Table1. Size, PDI and EE of screening lipids**

| Lipid | Size | PDI | EE |
| --- | --- | --- | --- |
| MC3 | 104.8 | 0.061 | 79.7% |
| A1-EPO15-O18B | 111.9 | 0.152 | 95.3% |
| A1-EP10A-O18B | 96.6 | 0.112 | 84.7% |
| A1-EP10-O18B | 105.1 | 0.119 | 93.9% |
| A1-EPO15-O18A | 104.6 | 0.148 | 92.3% |
| A1-EP10A-O18A | 102.9 | 0.045 | 89.0% |
| A1-EP10-O18A | 113.9 | 0.142 | 87.5% |

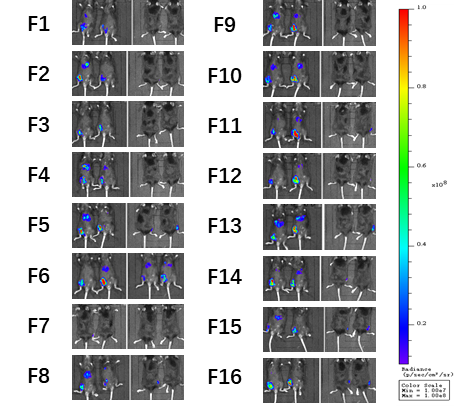

**Supplementary Figure 1.** **In vivo results of DOE study**

**Supplementary Table 2. Optimized formulation ratio of F11**

| **Compound** | **Lipid** | **DSPC** | **Chol** | **PEG** |
| --- | --- | --- | --- | --- |
| **Ratio(mol/mol)** | 46.0 | 10.0 | 42.4 | 1.6 |

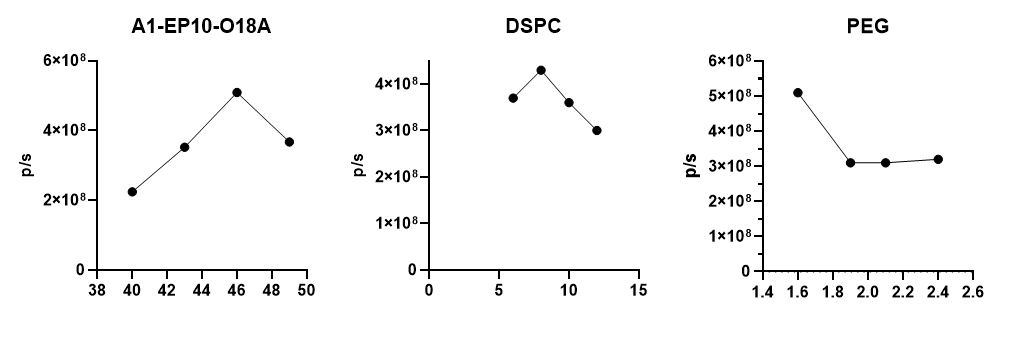

**Supplementary Figure 2. Analysis of In vivo results of DOE study**

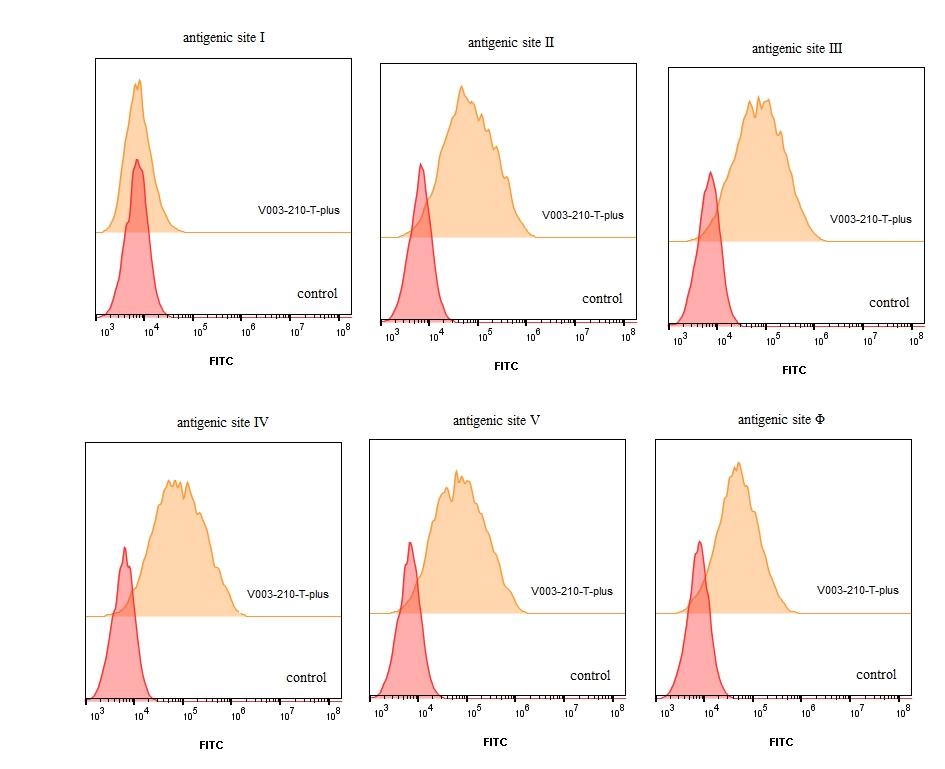

**Supplementary Figure 3. RSV F protein specific antigenic sites of V003-210-T-plus detected by FACS.** Antibodies recognizing antigenic site Ⅰ, Ⅱ, Ⅲ, Ⅳ, Ⅴ and Φ were used to stain RSV preF protein in HT1080 cells. Flow cytometric data were quantitatively evaluated using FlowJo software.

**Supplementary Table 2 Histopathology Semiquantitative Scoring Criteria**

| **Histological characterization** | **Score** | **Criteria** |
| --- | --- | --- |
| alveolar area  Inflammatory infiltration of macrophages, lymphocytes and neutrophils in alveolar tissue | 0 | none or no visible lesion |
|  | 1 | mild |
|  | 2 | moderate |
|  | 3 | marked |
|  | 4 | severe |
| Perivascular  Inflammatory infiltration of macrophages, lymphocytes and neutrophils around blood vessels | 0 | none or no visible lesion |
|  | 1 | mild |
|  | 2 | moderate |
|  | 3 | marked |
|  | 4 | severe |
| Peribronchial  Inflammatory infiltration of macrophages, lymphocytes and neutrophils around the bronchioles | 0 | none or no visible lesion |
|  | 1 | mild |
|  | 2 | moderate |
|  | 3 | marked |
|  | 4 | severe |
| Interstitial  Inflammatory infiltration of macrophages, lymphocytes and neutrophils in the interstitium | 0 | none or no visible lesion |
|  | 1 | mild |
|  | 2 | moderate |
|  | 3 | marked |
|  | 4 | severe |

**Supplementary Table 3 Grouping of Cotton Rats**

| Group | Gender | Immunization  Route | Immunizing Dose | Challenge Strain |
| --- | --- | --- | --- | --- |
| G1 | 3♂ | IM | STR-V003 (5 μg) | RSV A2 |
| G2 | 2♀ 1♂ | IM | STR-V003 (20 μg) | RSV A2 |
| G3 | 2♀ 1♂ | IM | Vehicle | RSV A2 |
| G4 | 1♀ 2♂ | IM | STR-V003 (5 μg) | RSV B9320 |
| G5 | 2♀ 1♂ | IM | STR-V003 (20 μg) | RSV B9320 |
| G6 | 2♀ 1♂ | IM | Vehicle | RSV B9320 |

**Supplementary Table 4 Pathology scores of lung tissues in cotton rats on Day 54**

| **Group** | **Histological characterization** | | | |
| --- | --- | --- | --- | --- |
|  | **Perivasculitis** | **Peribronchiolitis** | **Alveolitis** | **Interstitial pneumonia** |
| G1 STR-V003 5ug/animal (RSV A2) | 0 | 1 | 0 | 0 |
|  | 1 | 1 | 0 | 0 |
|  | 1 | 1 | 0 | 0 |
| G2 STR-V003 20ug/animal (RSV A2) | 0 | 1 | 0 | 1 |
|  | 0 | 1 | 0 | 0 |
|  | 0 | 1 | 0 | 0 |
| G3 Vehicle (RSV A2) | 0 | 0 | 0 | 0 |
|  | 1 | 1 | 0 | 0 |
|  | 0 | 1 | 0 | 0 |
| G4 STR-V003 5ug/animal (RSV B9320) | 0 | 2 | 0 | 0 |
|  | 0 | 2 | 0 | 0 |
|  | 0 | 2 | 0 | 0 |
| G5 STR-V003 20ug/animal (RSV B9320) | 0 | 1 | 0 | 0 |
|  | 0 | 1 | 0 | 0 |
|  | 0 | 2 | 0 | 0 |
| G6 Vehicle (RSV B9320) | 1 | 2 | 0 | 0 |
|  | 1 | 2 | 0 | 0 |
|  | 0 | 1 | 0 | 0 |

**Supplementary Table 5 Organ Weights Changes (Day 32)**

| Parameter  (% Change ^a^) | | Negative Control | STR-V003 (dose/animal) | | | Empty LNPs (dose/animal) | | |
| --- | --- | --- | --- | --- | --- | --- | --- | --- |
|  |  |  | 1 | 1.5 | 2 | 1 | 1.5 | 2 |
| **M** | **Thymus** | | | | | | | |
|  | Organ Weights (g) | 0.4916 | -35.9%* | -40.7%* | -52.4%* | -28.4%* | -41.9%* | -52.8%* |
|  | organ-to- brain weight ratios (%) | 23.2654 | -34.8%* | -37.6%* | -48.7%* | -25.7% | -39.7%* | -51.6%* |
|  | organ-to-body weight ratios (%) | 0.1102 | -31.3%* | -32.6%* | -45.7%* | -22.3% | -36.3%* | -46.7%* |
|  | **Spleen** | | | | | | | |
|  | Organ Weights (g) | 0.8326 | +15.6% | +24.2% | +31.9% | +10.4% | +15.0% | +18.3% |
|  | organ-to- brain weight ratios (%) | 39.5158 | +17.3% | +31.0%* | +42.1%* | +14.2% | +18.8% | +20.7% |
|  | organ-to-body weight ratios (%) | 0.1869 | +24.9% | +39.9%* | +51.4%* | +19.5% | +27.3% | +34.1%* |
| **F** | **Thymus** | | | | | | | |
|  | Organ Weights (g) | 0.4630 | -17.8% | -39.8%* | -42.8%* | -37.5%* | -36.2%* | -25.9%* |
|  | organ-to- brain weight ratios (%) | 23.8299 | -19.0% | -38.1%* | -42.1%* | -36.2%* | -33.5%* | -27.0%* |
|  | organ-to-body weight ratios (%) | 0.1608 | -13.7% | -34.8%* | -38.5%* | -32.1%* | -31.5%* | -23.9% |
|  | **Spleen** | | | | | | | |
|  | Organ Weights (g) | 0.6440 | +26.0%* | +22.3%* | +32.1%* | +17.8% | +18.9%* | +35.2%* |
|  | organ-to- brain weight ratios (%) | 33.1351 | +23.9%* | +25.6%* | +34.3%* | +20.1%* | +23.8%* | +33.5%* |
|  | organ-to-body weight ratios (%) | 0.2236 | +32.3%* | +32.9%* | +41.7%* | +28.8%* | +27.5%* | +38.8%* |

Note: "*" indicates *P*≤ 0.05 compared with the negative control group; "-" indicates decrease; "+" indicates increase; a represents % change compared with the negative control group. M = Male; F = Female.
